# Cooperative conformational transitions and the temperature dependence of enzyme catalysis

**DOI:** 10.1101/2023.07.06.548038

**Authors:** Emma J. Walker, Carlin J. Hamill, Rory Crean, Michael S. Connolly, Annmaree K. Warrender, Kirsty L. Kraakman, Erica J. Prentice, Alistair Steyn-Ross, Moira Steyn-Ross, Christopher R. Pudney, Marc W. van der Kamp, Louis A. Schipper, Adrian J. Mulholland, Vickery L. Arcus

**Author notes:** These authors contributed equally to the work (Emma J. Walker, Carlin J. Hamill).

## Abstract

Many enzymes display non-Arrhenius behaviour with curved Arrhenius plots in the absence of denaturation. There has been significant debate about the origin of this behaviour and recently the role of the activation heat capacity 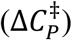 has been widely discussed. If enzyme-catalysed reactions occur with appreciable negative values of 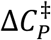 (arising from narrowing of the conformational space along the reaction coordinate), then curved Arrhenius plots are a consequence. To investigate these phenomena in detail, we have collected high precision temperature-rate data over a wide temperature interval for a model glycosidase enzyme MalL, and a series of mutants that change the temperature-dependence of the enzyme-catalysed rate. We use these data to test a range of models including macromolecular rate theory (MMRT) and an equilibrium model. In addition, we have performed extensive molecular dynamics (MD) simulations to characterise the conformational landscape traversed by MalL in the enzyme-substrate complex and an enzyme-transition state complex. We have crystallised the enzyme in a transition state-like conformation in the absence of a ligand and determined an X-ray crystal structure at very high resolution (1.10 Å). We show (using simulation) that this enzyme-transition state conformation has a more restricted conformational landscape than the wildtype enzyme. We coin the term “transition state-like conformation (TLC)” to apply to this state of the enzyme. Together, these results imply a cooperative conformational transition between an enzyme-substrate conformation (ES) and a transition-state-like conformation (TLC) that precedes the chemical step. We present a two-state model as an extension of MMRT (MMRT-2S) that describes the data along with a convenient approximation with linear temperature dependence of the activation heat capacity (MMRT-1L) that can be used where fewer data points are available. Our model rationalises disparate behaviour seen for MalL and a thermophilic alcohol dehydrogenase and is consistent with a raft of data for other enzymes. Our model can be used to characterise the conformational changes required for enzyme catalysis and provides insights into the role of cooperative conformational changes in transition state stabilisation that are accompanied by changes in heat capacity for the system along the reaction coordinate. TLCs are likely to be of wide importance in understanding the temperature dependence of enzyme activity, and other aspects of enzyme catalysis.

## Introduction

Scientific discourse on the temperature dependence of enzyme catalysis has a long history. In the last decade, debate has been a reignited regarding the origins of non-Arrhenius behaviour seen for many enzymes.^1–5^ We have argued previously that the changes in the conformational landscape along the reaction coordinate lead to changes in heat capacity that explain curved Arrhenius plots and we have given this scheme the name “macromolecular rate theory (MMRT)”.^6,7^ Others have used similar arguments of conformational complexity to explain deviations from Arrhenius behaviour and we have suggested that these proposals are complementary.^5,6,8^

Conformational changes can be envisaged as ‘many-state’ or ‘two-state’, where the latter often implies cooperativity for macromolecules. Cooperative phenomena are ubiquitous in biology and are a feature of protein folding, ligand binding, enzyme catalysis and allosteric regulation.^9^ Two-state protein folding is an archetype for cooperativity, with an equilibrium between a unfolded state and a folded state involving large numbers of intramolecular interactions. The observed folding kinetics for two-state protein folding are the sum of the forward and reverse rate constants, and the steady state equilibrium is the quotient of these two rate constants at a given temperature.^9^ The cooperativity of the transition leads to a change in heat capacity between the two states and a curved temperature-dependence of the equilibrium free energy (Δ*G*) giving rise to denaturation at both low and high temperatures.^10,11^ The position of the transition state for folding with respect to the folded and unfolded states also dictates the temperature-dependence of the activation free energy (Δ*G*^‡^) and hence the temperature-dependence of the folding and unfolding kinetics.^12^ For example, for barnase, the temperature dependence of the folding kinetics deviates significantly from Arrhenius kinetics due to the transition state for folding closely resembling the folded state, giving rise to a significant value of the activation heat capacity 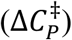 for folding 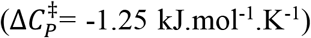.^12^ In contrast, the unfolding kinetics are close to Arrhenius in behaviour for the same reason – the transition state closely resembles the folded state and hence a negligible value of 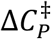 upon unfolding. Importantly, the temperature dependence of the folding kinetics for barnase gives rise to positive activation enthalpies at low temperatures and negative activation enthalpies at high temperatures due to the steep temperature dependence of Δ*H*^‡^ (a consequence of 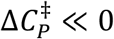). This is a hallmark of the kinetics of cooperative conformational changes involving changes in heat capacity along the reaction coordinate.

Large negative values for Δ*C*_*P*_ are seen for the binding of transition-state analogues to the enzyme 5’-methylthioadenosine phosphorylase (MTAP).^13^ This results in positive enthalpies for binding at low temperatures and negative enthalpies for binding at high temperatures due to the steep temperature dependence of Δ*H* for binding. The impressively small equilibrium binding constants for this system (*K*_*d*_ < 1 nM) highlight the very essence of enzyme catalysis, namely, stabilisation of the transition state species by the enzyme. The cooperative nature of this tight binding is manifested in the large, negative value for Δ*C*_*P*_ for this interaction (∼−2.3 *kJ. mol*^−1^. *K*^−1^). This equilibrium Δ*C*_*P*_ is reflected in the temperature dependence of the rate with a commensurate value of the activation heat capacity for MTAP with 2-amino-5’-methylthioadenosine as substrate 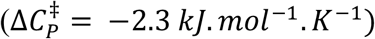.^13^

The importance of activation heat capacity for enzyme kinetics has been the subject of debate recently.^3,5,6,14^ The origin of this phenomenon lies in differences in fluctuations and populations of conformations between the enzyme-substrate complex (ES) and the enzyme-transition state complex (E-TS). We have argued that a consensus is emerging whereby the enzyme-substrate complex fluctuates between at least two conformations, one that favours the substrate and one that favours the transition state and that there is a difference in heat capacity between these two conformations.^6^ Arguments for an equilibrium between two states in the context of enzyme kinetics date back at least 70 years^1^ and also underpins the MWC and KNF models for allostery.^15^ Cooper and Dryden proposed (37 years ago) increased fluctuations about a mean conformation as an equivalent mechanism underlying allostery^16^ and indeed, an ensemble view of allostery has recently gained attention^17^ and acceptance.^18^ More recently, it has been shown that apparently puzzling temperature dependence of kinetic isotope effects in enzyme-catalysed reaction can be accounted for by a transition state theory model including two states with different reactivity.^19^ It can be argued that all these mechanisms involving multiple conformational states, coupled with changes to the number of states along the pathway, can give rise to a change in heat capacity if they involve changes in conformational fluctuations along the pathway.

Many investigators have invoked multiple conformations in the ES complex to explain deviations from Arrhenius kinetics for enzyme catalysed reactions. Truhlar and Kohen postulated an equilibrium between reactive and non-reactive states as an underlying mechanism giving rise to these deviations and noted similar behaviour for non-biochemical systems.^4^ This had been previously suggested by Massey and colleagues in 1966 for an amino acid oxidase.^20^ Daniel and Danson applied an “equilibrium” model (involving active and inactive states, ES and ES’) to explain deviations from Arrhenius kinetics for a wide range of enzymes.^21^ Mulholland and colleagues have also invoked two reactive states in enzyme catalysis and use transition state theory to rationalise deviations from Arrhenius kinetics.^19^ Åqvist and colleagues have made an effort to differentiate between an “equilibrium” model (with active and inactive states) and an activation heat capacity model (MMRT) proposed by us (also invoking multiple conformations).^5^ Although Åqvist presents these two models in opposition, we have argued that they are actually consistent, and effectively two sides of the same coin, whereby a constriction of the conformational space along the reaction coordinate (i.e. a two-conformation to one-conformation transition from ES to E-TS) can give rise to an appreciable activation heat capacity.^6^ More recently, Åqvist has used molecular simulations to argue that there is no evidence for 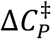 in enzyme-catalysed reactions.^22^ In contrast, Warshel and colleagues have used extensive MD/EVB simulations to show temperature-dependent activation enthalpies and entropies for alcohol dehydrogenase (ADH) and thus, by definition, an activation heat capacity for this enzyme.^8^ They argue that pre-organisation is generally important for enzyme catalysis.^8^

Here, we analyse these questions in detail through a combination of structural and biochemical experiments, molecular simulations and kinetic modelling on a well characterized enzyme, which has been the focus of debates and proposals in this area. We present high-resolution temperature-rate data for the model glycosidase enzyme MalL (over a wide temperature range) and for a range of mutants that alter the temperature-dependence of the enzyme catalysed reaction rate. We combine these kinetic data with detailed molecular dynamics simulations, and high-resolution X-ray crystal structures where different conformations are trapped. Our MD simulations show that the ES complex accesses a range of conformations whereas the E-TS complex is much more constrained. We call this conformation the transition-state-like conformation (TLC). This is a conformation that favours the chemical transition state and is visited by the enzyme when the substrate is bound and is the conformational bottleneck in phase space before the chemical step. For MalL, this is a cooperative transition that involves the shortening of more than 28 hydrogen bonds, an increase in correlated motions, and thus a significant increase in order immediately prior to the chemical step. This order-disorder transition (the ES-TLC transition) is temperature dependent, and we argue that this is the origin of the activation heat capacity.

We present and test alternative models to describe the temperature dependence of reaction for MalL including: a two-state model that is an extension of MMRT (MMRT-2S); and a convenient function with linear temperature-dependence for 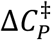 that serves as a good approximation to describe the kinetic data (MMRT-1L). We also apply this model to the intriguing case of alcohol dehydrogenase, where we show that the temperature dependent transition between ES and TLC is reversed (compared to MalL) due to the ES complex being more ordered than the TLC complex leading to the opposite slope for 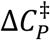. We show how our analysis for the temperature dependence of Δ*H*^‡^ and Δ*S*^‡^ for alcohol dehydrogenase is in agreement with the results of Warshel and colleagues, emphasising the emerging consensus.^8^

## Results

### Temperature-dependent enzyme kinetics and analysis

We collected high-quality enzyme kinetics data over a wide temperature range (6-56 °C), giving a dataset that allows a thorough analysis of the temperature dependence of MalL and its mutants. We extensively optimised our stopped-flow experimental protocols to ensure highly reproducible data and precise temperature control. A feature of the optimised protocol is the use of several dummy shots before each data collection run to ensure a very stable temperature throughout the stopped flow lines. The temperature-dependence data for WT MalL (in the absence of denaturation), collected at pH 7.0, are shown in Figure 1A&C. We have plotted all three replicates: note how tightly constrained the errors are for each point. We use transition state theory (equation 1) with a transmission coefficient (γ) of 1 to convert temperature-rate data (Figure 1C) into the activation free energy at a given temperature (Figure 1A). The temperature-dependence of Δ*G*^‡^ shows two regimes, with a clear transition at ∼313 K. There are sufficient data to initially treat these regimes separately. Each shows curvature (non-Arrhenius behaviour) and therefore MMRT fitting is appropriate (using equation 2).^7^ This gives 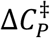 values of ∼0.8 (i.e. near zero) and ∼^−^20.8 kJ.mol^-1^.K^-1^ for the low and high temperature regimes respectively (blue and red curves Figure 1A). This immediately suggests that 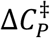 is temperature dependent and that there is a two-state cooperative transition at ∼313 K (Figure 1B).

**Figure 1.**
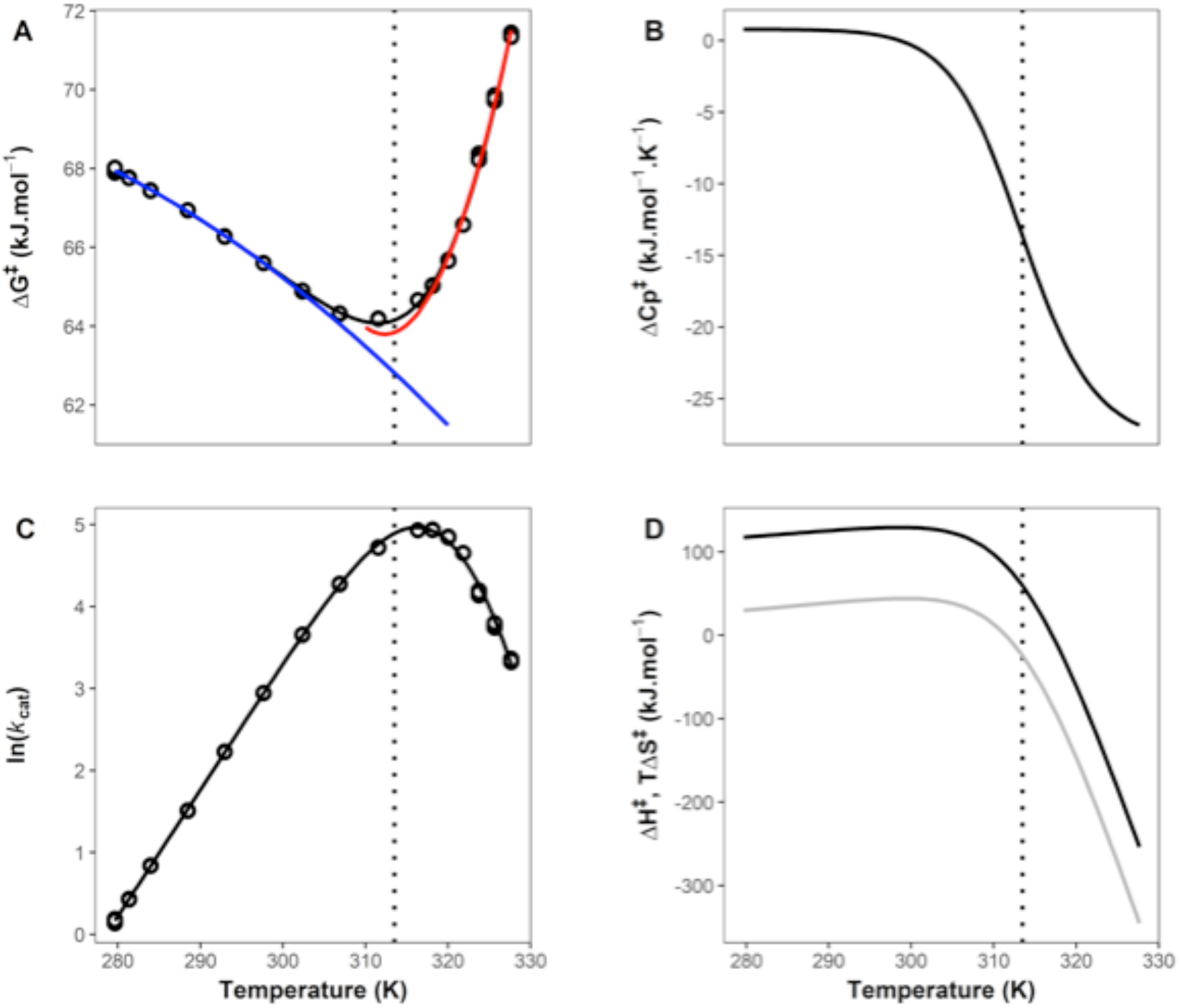
High-resolution temperature dependence of activity for MalL. **A**. Δ*G*^‡^ versus temperature. All data are plotted as circles (triplicates at each temperature). The blue curve is a plot using MMRT^7^ (equation 2) for the first six temperature points; the red curve is an MMRT fit for the last six temperature points. The black curve in **A** uses the two-state model proposed here (MMRT-2S), which shows a smooth transition between the blue and red curves (equations 4 & 5). **B**. 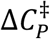 versus *T* illustrating the two-state model with constant 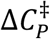 values at low and high temperatures and a cooperative transition between the two. **C**. ln *k*_*cat*_ versus *T*. Experimental data points in triplicate are plotted as circles. The curve is fitted to the two-state model (equations 4 & 5). **D**. Derived values for Δ*H*^‡^(black) and *T*Δ*S*^‡^ (grey) versus *T*. The transition between low and high temperature regimes is clear. In all panels, the vertical dotted line shows the transition temperature, *T*_*C*_ (313.5 K, determined from fitting equations 4 & 5).

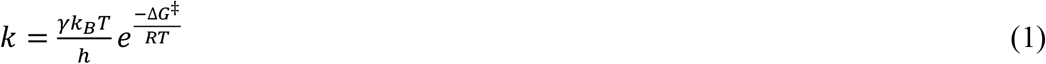

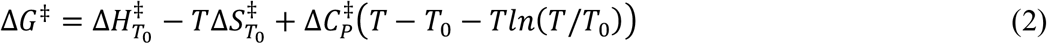

To establish a model to describe these data, we begin with two conformations: the enzyme-substrate conformational ensemble (ES) and a transition-state-like conformation (TLC) that favours the chemical transition state and the chemical reaction. In this context we differentiate between TLC and E-TS to highlight the fact that the TLC state is a conformational state visited from the ES complex and these conformations (ES and TLC) are on pathway. For the E-TS state in MalL in crystallography and in MD simulations, we use the complex of the enzyme with a transition-state analogue.

A minimal model is:

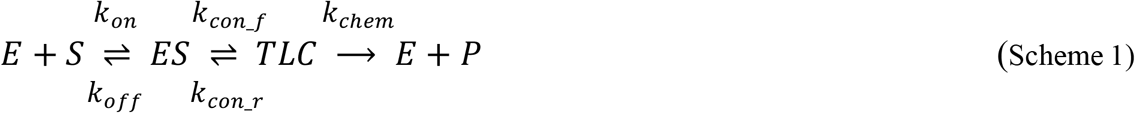

The substrate binding equilibrium is followed by a conformational equilibrium between ES and TLC and the chemical step proceeds from the TLC. The rate constants in Scheme 1 are for forward and reverse binding of the substrate (*k*_*on*_, *k*_*off*_), the forward and reverse conformational transitions between ES and TLC (*k*_*con_f*_, *k*_*con_r*_) and the rate constant for the chemical step (*k*_*chem*_). Formally, the rate equation for this system is:

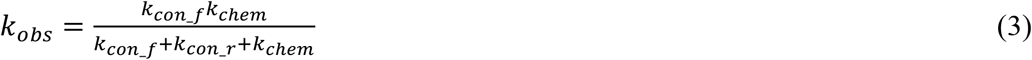

However, this is unwieldy as each rate constant potentially has three variables (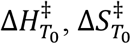 and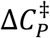, equations 1 & 2) giving an expression with potentially nine variables. We therefore take a simplified approach.

The two arms of the rate and activation free energy data suggest a two-state transition between low temperatures and high temperatures with a transition at ∼313 K. The value of 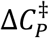 at low temperatures is near zero, while the value at high temperatures is large and negative. We can construct a suitable model for such a two-state transition by treating the 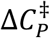 as a function of temperature thus:

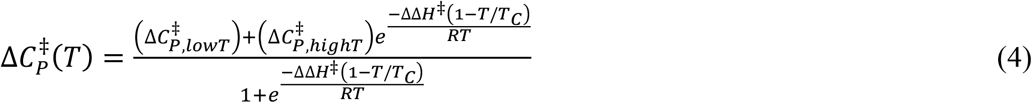

…where *T*_*C*_ is the temperature at the midpoint of the transition, 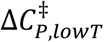 is the value of 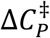, for the low temperature arm, 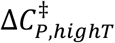 is the value of 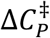 for the high temperature arm and ΔΔ*H*^‡^ is the difference in Δ*H*^‡^ between the two arms (at *T*_*C*_). Having defined 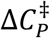 as a function of temperature, the activation enthalpy, entropy and free energy terms are given by standard expressions:

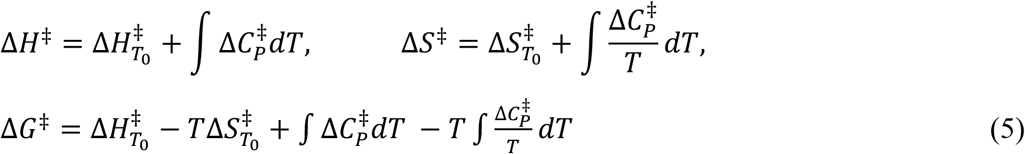

The rate coefficient at a particular temperature is given by equation 1. This provides a rate equation with five terms, 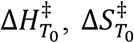, *T*_*C*_, ΔΔ*H*^‡^ and 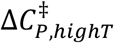 with 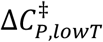 having been found first from an independent fit of the low temperature arm of the data (blue curve in figure 1A).

We have collected very accurate kinetic data for WT MalL and for the mutant series V200T, V200S and V200A and fitted these data using our proposed two-state model, MMRT-2S (Figure 2 and Table 1). The data are very well described by this model, and in all cases show a transition between 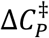 near zero for low temperatures and large negative values for 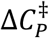 at high temperatures (Figures 1B&2B). The values for Δ*C*^‡^ at high temperatures for the mutant series get less negative along the series (WT-V200A-V200T-V200S) which is consistent with these mutations increasing the rigidity of the ES complex as we have described previously^3,7^ (Figure 2B). The value found for 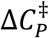 at *T*_*C*_ is also consistent with previous results measured over a narrow temperature range around 310 K.^3^ The significant increase in rate for V200A comes at the cost of *K*_*M*_ for this mutation: the *K*_*M*_ value is more than 7-times that of the WT (*K*_*M*_ = 1.56 mM, c.f. 0.21 mM for WT).^3^ Mutation from valine at this position in all cases significantly changes the temperature dependence of the rate and shifts *T*_*opt*_ up by ∼6 degrees. It is interesting to note that the WT enzyme, which has a more flexibile ES state, has a slightly higher rate at intermediate temperatures (c.f. V200T and V200S): this is consistent with previous hypotheses regarding the evolution to psychrophily (cold adapted activity) whereby higher flexibility is observed for psychrophilic enzymes than their mesophilic counterparts.^7,23,24^ This increased flexibility of the ES comes at the cost of a lower *T*_*C*_ and a larger, more negative 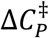 consistent with the ‘psychrophilic trap’ that we have previously described.^7^ The steepness of the transition from low temperature to high temperature is determined by the difference in Δ*H*^‡^ between the two processes (ΔΔ*H*^‡^) (although this parameter is not particularly well determined by the data: relatively large standard errors of fitting, see Table 1). As expected ΔΔ*H*^‡^ is significantly lower for V200S than for the WT: the V200S ES complex is more rigid.

**Table 1.**
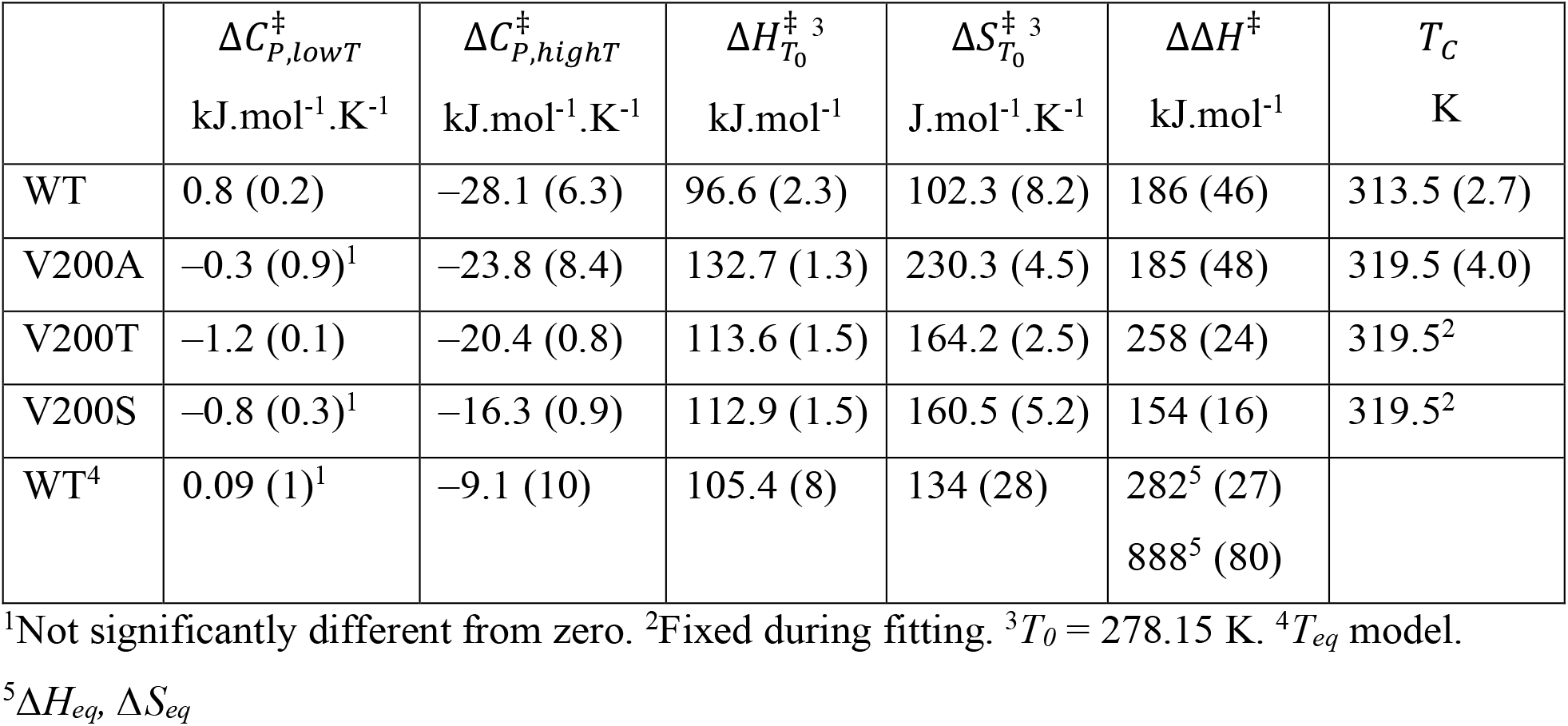
Activation parameters from the fitting to the two-state model (MMRT-2S).

| | $\Delta C_{P,lowT}^\ddagger$<br>kJ.mol<sup>-1</sup>.K<sup>-1</sup> | $\Delta C_{P,highT}^\ddagger$<br>kJ.mol<sup>-1</sup>.K<sup>-1</sup> | $\Delta H_{T_0}^\ddagger$<sup>3</sup><br>kJ.mol<sup>-1</sup> | $\Delta S_{T_0}^\ddagger$<sup>3</sup><br>J.mol<sup>-1</sup>.K<sup>-1</sup> | $\Delta\Delta H^\ddagger$<br>kJ.mol<sup>-1</sup> | $T_C$<br>K |
| --- | --- | --- | --- | --- | --- | --- |
| WT | 0.8 (0.2) | -28.1 (6.3) | 96.6 (2.3) | 102.3 (8.2) | 186 (46) | 313.5 (2.7) |
| V200A | -0.3 (0.9)<sup>1</sup> | -23.8 (8.4) | 132.7 (1.3) | 230.3 (4.5) | 185 (48) | 319.5 (4.0) |
| V200T | -1.2 (0.1) | -20.4 (0.8) | 113.6 (1.5) | 164.2 (2.5) | 258 (24) | 319.5<sup>2</sup> |
| V200S | -0.8 (0.3)<sup>1</sup> | -16.3 (0.9) | 112.9 (1.5) | 160.5 (5.2) | 154 (16) | 319.5<sup>2</sup> |
| WT<sup>4</sup> | 0.09 (1)<sup>1</sup> | -9.1 (10) | 105.4 (8) | 134 (28) | 282<sup>5</sup> (27)<br>888<sup>5</sup> (80) |  |

<sup>1</sup>Not significantly different from zero. <sup>2</sup>Fixed during fitting. <sup>3</sup> $T_0 = 278.15$ K. <sup>4</sup> $T_{eq}$ model.
<sup>5</sup> $\Delta H_{eq}$ , $\Delta S_{eq}$

**Figure 2.**
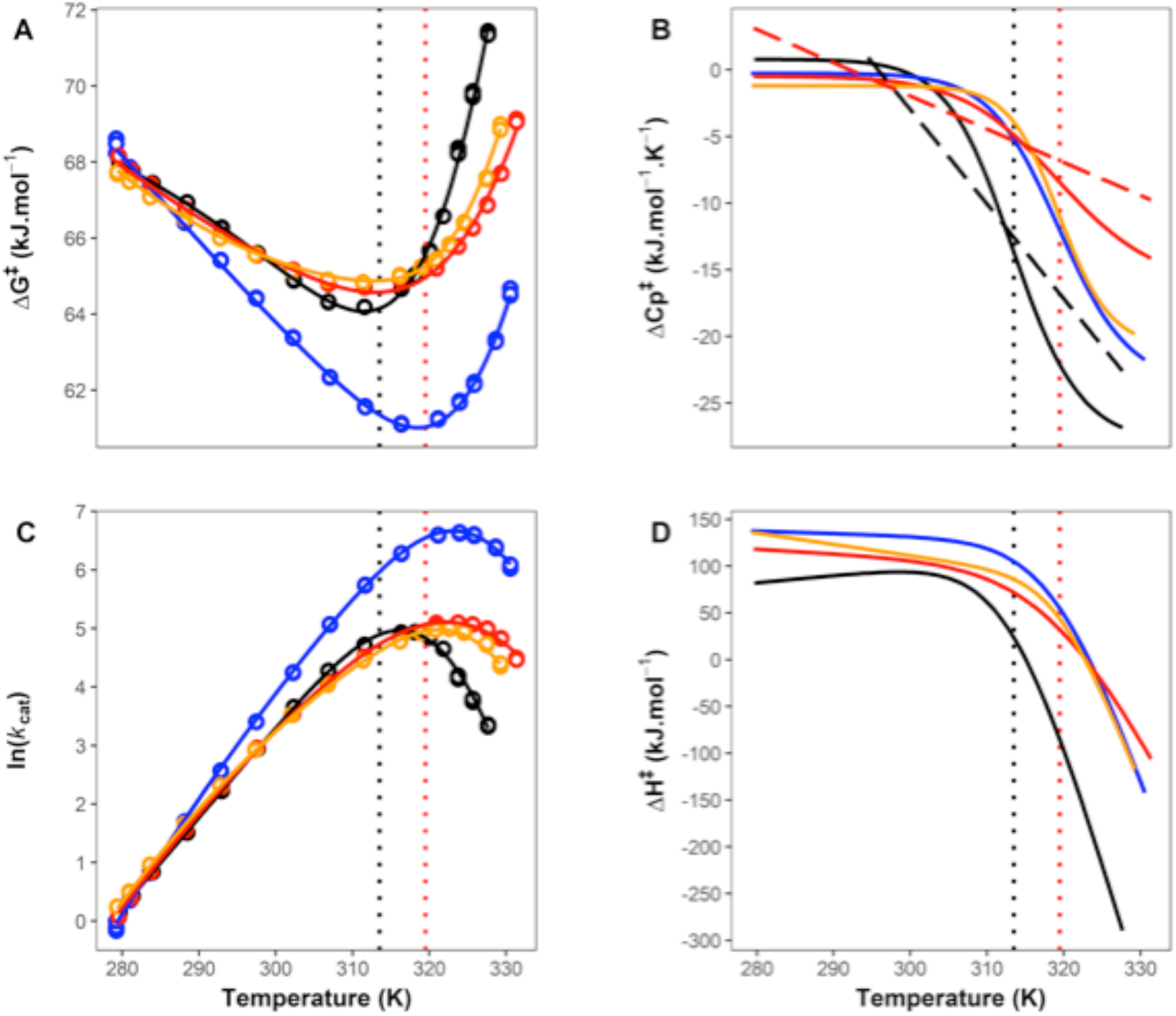
Temperature-dependent enzyme kinetic measurements for MalL WT (black), V200T (orange), V200S (red) and V200A (blue) mutations. **A**. Δ*G*^‡^ versus T. **B**. 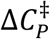 versus *T*. **C**. ln(*k*_*cat*_) versus T. **D**. Δ*H*^‡^ versus T. All smooth curves are fits of MMRT-2S (equations 4 & 5) to the data and in the case of 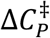 and Δ*H*^‡^, are derived from these fits (see Table 1). Vertical dotted lines are values for *T*_*C*_ for WT (black) and the V200T, V200S and V200A mutants (red; all three have the same *T*_*C*_ values). Linear dashed lines (panel B) are linear 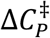 values from fitting the data using MMRT-1L (equations 6 & 1) for WT and V200S.

Figure 4D clearly illustrates the pitfalls of discussions based on activation enthalpies and entropies for enzyme-catalysed reactions: activation enthalpy curves for WT MalL and the V200 variants cross, with the order at low temperatures being reversed at moderate temperatures and changing significantly again at high temperatures. Discussions of enzyme activity, and e.g. of the evolution of enzyme catalysis, must consider this temperature dependence; arguments based on enthalpy or entropy are not independent of temperature. Notwithstanding this, large positive values of Δ*H*^‡^ at low temperatures are consistent with a bond-breaking chemical step and large negative values of Δ*H*^‡^ at high temperatures suggest that a cooperative conformational step contributes to the observed rate with formation of hydrogen bonds along the reaction coordinate. In chemical systems, negative activation enthalpies have been attributed to the formation of multiple hydrogen bonds *en route* to the transition state.^25^

### A convenient approximation, with linear temperature dependence of 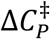 (MMRT-1L)

The two-state model presented above requires fitting of five or six parameters; at least two of the parameters are tightly correlated giving rise to practical issues with the fitting procedure (e.g. parameters not converging). For example, in the case of V200T and V200S, the value of *T*_*C*_ must be fixed to prevent this from happening. Indeed, most enzyme rate-temperature datasets contain fewer data points and thus cannot be reasonably fitted using a five-parameter model. Hence, a simpler model that captures the trends in the data is warranted. We propose a linear temperature-dependence of 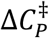 as a useful approximation to MMRT-2S, because the slope of 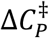 captures the dynamics of the transition. For example, the slope of 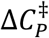 for such a linear model would encompass both the size of 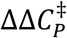 across the temperature range and the steepness of the transition as reflected in the size of ΔΔ*H*^‡^. For a such a linear model, we take 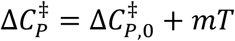, where 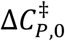 is the y-axis intercept at 0 K and *m* is the slope of 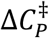. We derive a function for Δ*G*^‡^ by integration in the usual way (equation 5). This results in an additional *T*^*2*^ term in the activation free energy (equation 6):

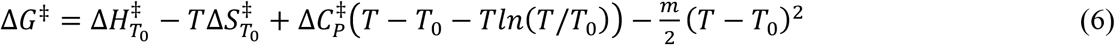

The enthalpy and entropy are given by:

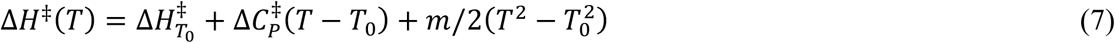

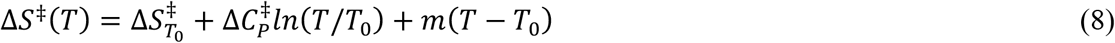

In the context of the rate equation, the maximum temperature for reaction (*T*_*opt*_) and the inflection points (*T*_*inf*_)^26^ are:

*T*_*opt*_ *when* Δ*H*^‡^ = −*RT*

*T*_*inf*_ *when* 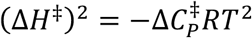 i.e. the criteria for the variance where 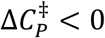.

Here, we have used MMRT-1L to fit the rate data for WT and V200S MalL. The linear temperature dependence of 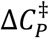 is shown in Figure 2B (black and red dashed lines and shows slopes of –710 and –252 J.mol^-1^.K^-2^ respectively). These lines capture the trend for the two-state transition and their slopes capture both the magnitude of the change in 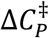 and the steepness of the transition (ΔΔ*H*^‡^).

As an example of the utility of MMRT-1L, we take data from the literature for alcohol dehydrogenase (ADH) which has been extensively analysed by Klinmann *et al*. and Warshel *et al*. but has relatively few temperature points (7-8 points).^8,19,27,28^ Fitting equation 6 to these data for the protiated and deuterated substrates makes it immediately obvious that the temperature dependence of 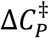 is opposite to that seen for MalL: for ADH, at low temperatures 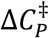 is negative and at higher temperatures it approaches zero (Figure 3). This is consistent with Warshel *et al*.’s analysis and hypothesis^8^ that the ES state is more ordered than (what we denote here as) the TLC state for ADH. Thus, the ES state is favoured at low temperatures, accompanied by negative values for 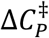 (in contrast to MalL where the TLC state is favoured at low temperatures; 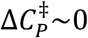). The cooperative transition between ES and TLC that we invoke is also consistent with the details of the differences in the temperature dependence of the rate between the protiated and deuterated substrates (over and above zero-point energy differences) where small changes can manifest as large differences in cooperative systems. We have previously observed differences of this magnitude for thermophilic glucose dehydrogenase with protiated and deuterated substrates.^29^ Warshel and colleagues highlight this point when they suggest a “phase transition” for the ADH system.^8^ We illustrate this ‘phase transition’ for ADH data by estimating the two-state transition for 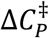 in Figure 3B (dashed lines). Here, at low temperatures, our model indicates that a cooperative conformational transition contributes to the free energy barrier and can be seen in large negative values for 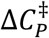. At high temperatures, the TLC state is favoured and the free energy barrier is dominated by the chemical step, with 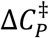 close to zero. Our analysis closely reproduces the analysis of Warshel and colleagues albeit via a completely different route and the two-state transition shows steep temperature-dependence of Δ*H*^*‡*^ and Δ*S*^*‡*^ below the *T*_*C*_ value of ∼305 K and constant values of Δ*H*^*‡*^ and Δ*S*^*‡*^ above the *T*_*C*_ with Δ*S*^*‡*^ close to zero at these higher temperatures (Figure 3D).

**Figure 3.**
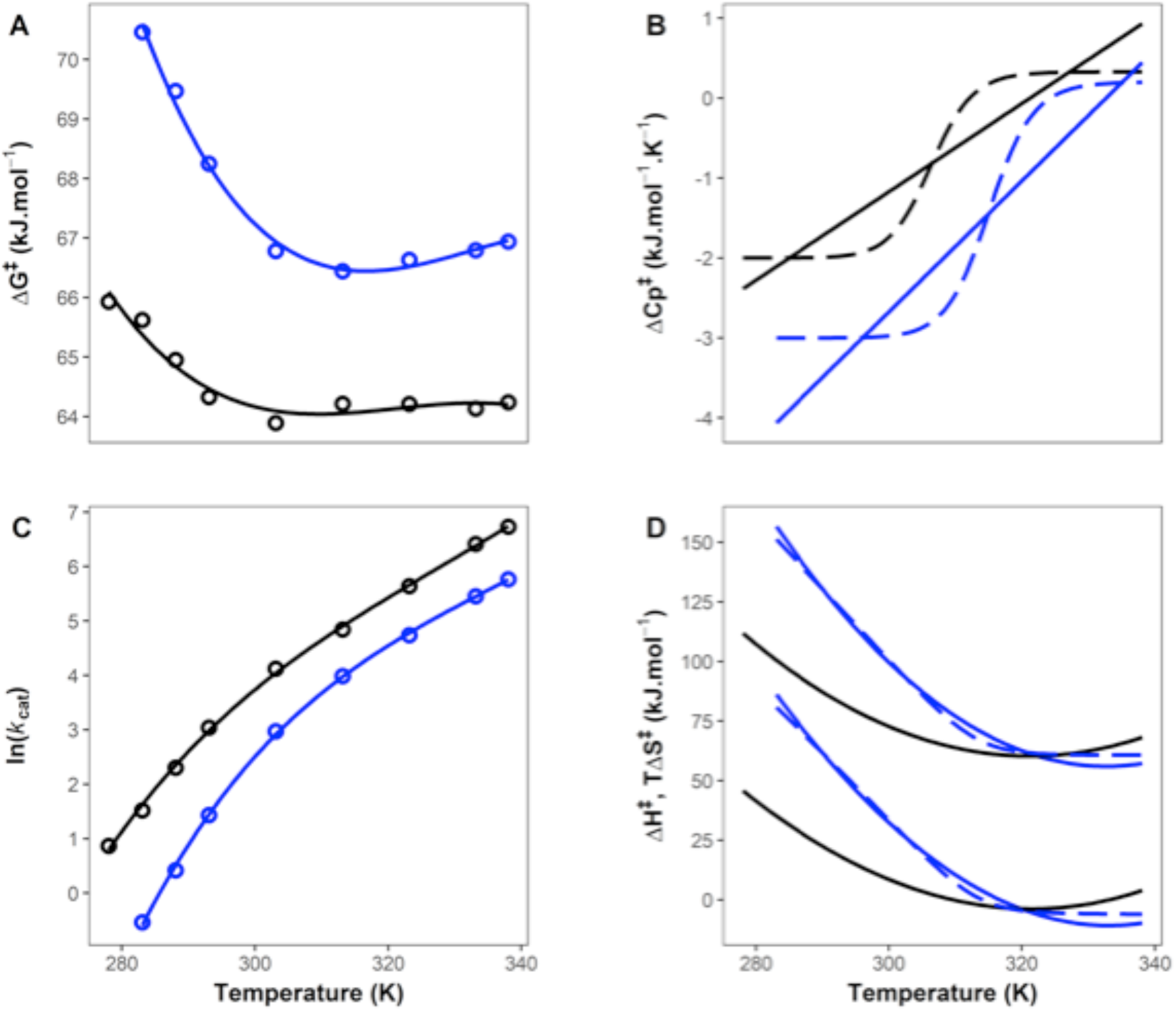
Temperature dependence of ADH for protiated and deuterated substrates. **A**. Δ*G*^‡^ versus T for protiated (black) and deuterated (blue) substrates. **B**. Derived values of 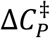 from fitting the linear model (MMRT-1L) to the data (equation 6).^28^ Two-state curves are shown as dashed lines in panel B and are for illustration as there are insufficient data to fit the two-state model (MMRT-2S). **C**. ln(*k*_*cat*_) versus T for protiated and deuterated substrates. **D**. Derived values of Δ*H*^*‡*^ and Δ*S*^*‡*^ for protiated and deuterated substrates (equations 7&8). The dashed lines are for the two-state model for the deuterated substrate by way of illustration.

### Two conformations on the pathway for the enzyme-catalysed reaction

Previously, we have used extensive molecular dynamics simulations to characterise the conformational landscape in two states (ES and E-TS) for the model enzymes ketosteroid isomerase (KSI) and MalL and designer Kemp eliminase enzymes including an evolved variant.^14,30^ This involved repeated independent simulations (x10) of 500 ns each to characterise the enzyme bound to either the substrate (ES complex) or a transition state analogue (E-TS complex). The difference in heat capacity between these states was calculated from the difference in fluctuations in the enthalpy for the two conformational ensembles. While calculation of heat capacities from this fluctuation approach has recently been criticised, we note that it has been applied successfully e.g. in the context of protein folding simulations^31^ and, for KSI and MalL, it gives results in very good agreement with experimental observations from MMRT fitting.

Here, we have applied these simulation and analysis approaches to characterise WT MalL in detail and to compare it with mutant MalL (see below). This involved 20 independent simulations of 500 ns each for both the ES complex and the E-TS complex. The latter is simulated by binding a (physical) transition-state analogue.^14^ Simulations were also performed for the mutant MalL S536R (see below). Principal component analysis was used to characterise the conformational landscape in each of the two states (Figure 4). These simulations show that the ES complex has a much broader conformational landscape than the E-TS complex and that the ES complex visits the E-TS conformation in traversing the landscape (Figure 4C&D). The E-TS complex is characterised by a constrained conformation with a significant number of additional H-bonds clustered in the centre of the structure when compared to the ES complex. The first principal component (PC1) for the ES complex is characterised by translational movement of the lid domains above the active site of the enzyme (Figure 4A&B). These movements are large and can be broadly captured by the Cα-Cα distance between His217 in domain 2 and Glu400 in domain 3 on opposite sides of the active site. This distance fluctuates between 35 Å and 15 Å and clusters around 25 Å and 17.5 Å corresponding to the two peaks in Figure 4C at -18 Å and 16 Å respectively. Thus, broadly speaking, the ES state fluctuates between “open” and “closed” conformations with domain movements of up to 20 Å. The E-TS complex corresponds to the “closed” conformation and our hypothesis is that the transition between open and closed is cooperative involving the establishment of a large number of intramolecular interactions (e.g. ∼28 shortened H-bonds, see below). This emphasises the extraordinary preorganisation necessary to achieve a rate enhancement of ∼10^15^ (i.e. a stabilisation of the transition state of ∼110 kJ.mol^-1^). The second principal component (PC2) is characterised by an opening and closing of the loops of domains 1 and 2 with the lid domain of domain 3 rotating as it closes (Figure 4B). The PC2 histogram shows that both states fluctuate around the same mean position and again, that the E-TS complex traverses a narrower conformational space when compared to the ES complex (see supplementary Figure 1). The two-dimensional plot of PC1 versus PC2 clearly shows the additional conformational space traversed by the ES complex when compared to the E-TS complex (Figure 4D). The origin of the heat capacity difference between ES and E-TS lies in the broad conformational fluctuations for the ES complex (primarily in lid and loop domains above the active site) in contrast to the narrowed fluctuations for the E-TS complex. Importantly, the E-TS conformation is visited by the ES conformations and formation of the E-TS complex is accompanied by the strengthening of a large number of hydrogen bonds. Indeed, these observations are commensurate with the very tight binding of the transition state chemical species by the enzyme which is important for catalysis.

**Figure 4.**
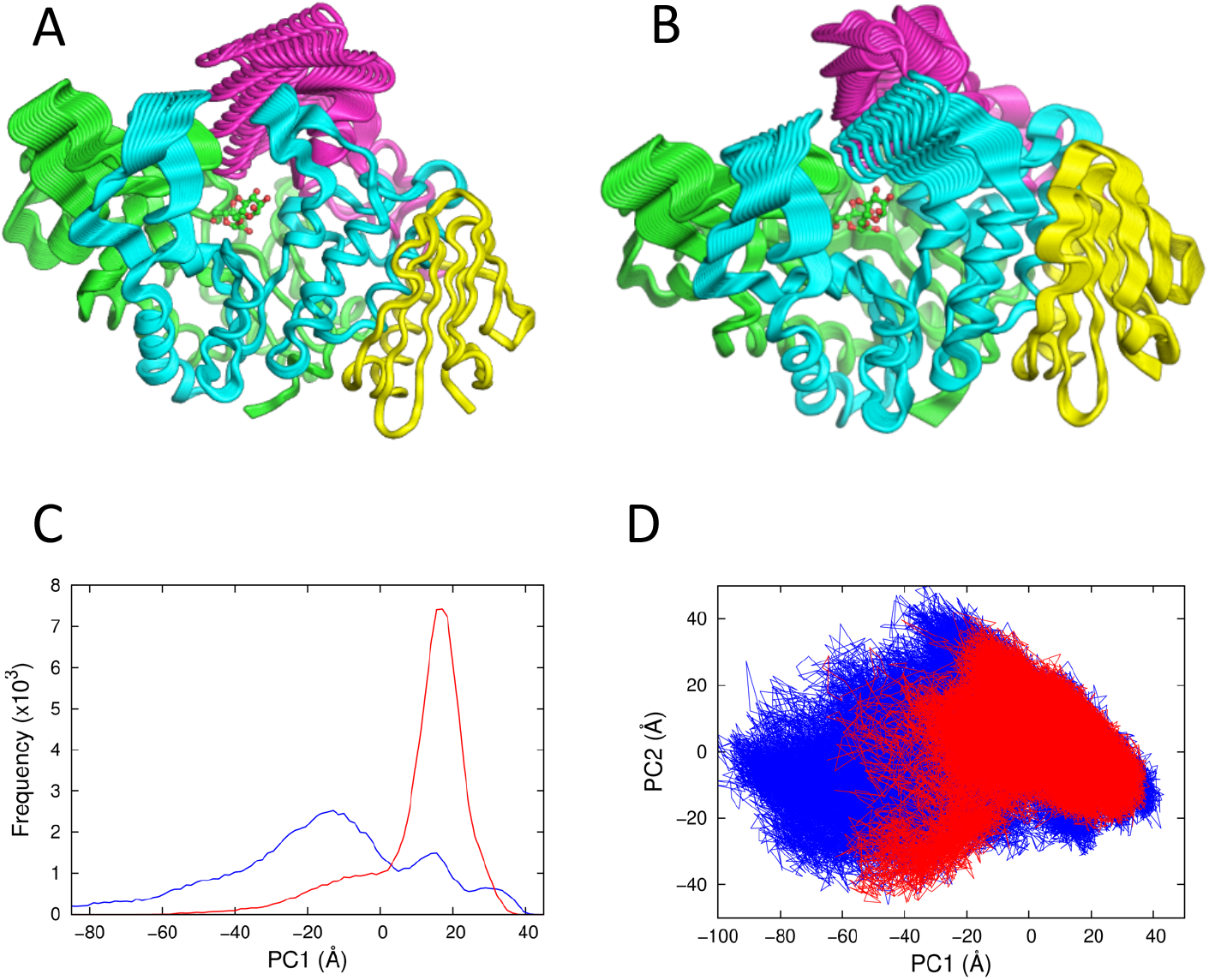
Conformational landscapes of WT MalL ES and E-TS complexes from MD simulations. **A**. Structure of MalL WT showing the first principal component (PC1) projections illustrating the movement of domains for the ES complex. The structure is coloured to indicate 4 regions (1-193, green; 194-321, blue; 322-459 magenta; 460-561, yellow). The substrate (isomaltose) is shown as ball-and-stick in green and red. **B**. Principal component projections illustrating the movement of domains for the ES complex for PC2. **C**. Principal component analysis showing PC1 histogram for the ES complex (blue) and E-TS complex (red). **D**. Two-dimensional plot of PC1 versus PC2 for the ES complex (blue) and the E-TS complex (red).

### Very high resolution structure of apo-S536R that favours the TLC

We designed several mutants based on WT MalL X-ray structures determined in the presence of urea. We designed arginine mutations to substitute the guanidinium group of the arginine side chain for urea thereby introducing new hydrogen bond networks at the surface of the protein. Our aim was to assess the allosteric effects of these new interactions on the enzymatic rate and conformational dynamics. In the case of S536R we determined a structure at very high resolution in the absence of ligand (1.10 Å resolution, Supplementary Table 1, PDB code 7LV6). Notably, there are only very small differences between the temperature-dependent enzymatic activity of the mutant enzyme (S536R) and WT MalL (Supplementary Figure 2).

Although the overall RMSD between the WT and S536R crystal structures is small (0.185 Å over 496 Cα positions), this masks very significant differences in the H-bonding patterns for the two conformations. The structures show significant shortening (>0.3 Å) of 28 H-bonds in the S536R structure compared to the WT structure, which is consistent with increased order of the mutant structure (Figure 5, for H-bond criteria and bond lengths see Methods section). The majority of the shortened H-bonds (20 H-bonds) are found in two regions of the enzyme: region 1, 1-193 (green, Figure 5A) and region 3, 322-459 (magenta, Figure 5A). Further, an additional H-bond between the mainchain of Tyr14 and the sidechain of Gln369 connects these two regions. These two regions are contiguous in the structure and are the most mobile in the MD simulations for the WT enzyme discussed above. Their reduced mobility in the E-TS complex makes the greatest contribution to the calculated 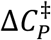 from the WT simulations.^14^

**Figure 5.**
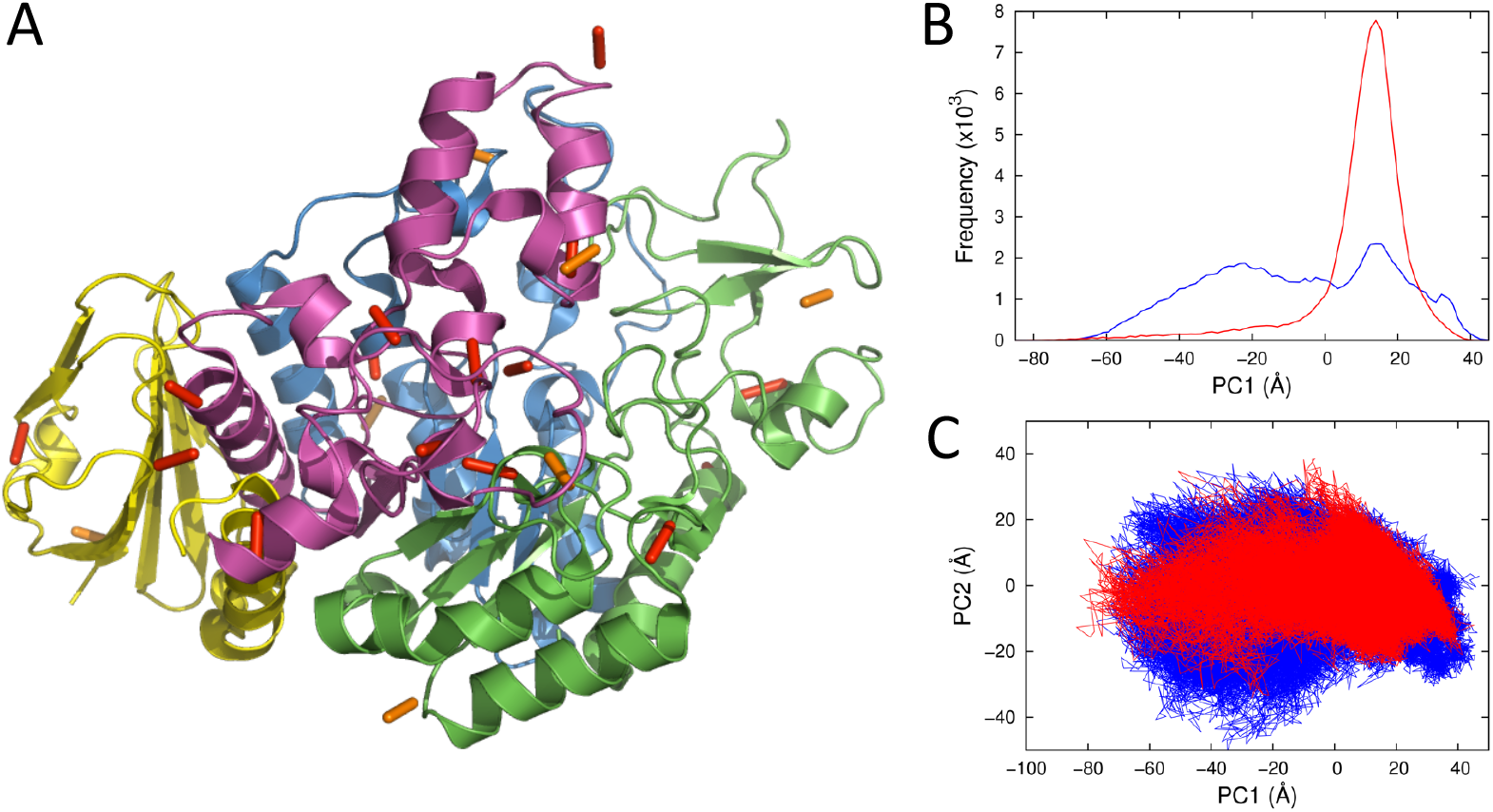
MalL S536R crystal structure and principal component analyses. **A**. The structure of MalL S536R determined at 1.10 Å resolution. The view is from the back (c.f. Figure 4) and regions are coloured the same as Figure 4. H-bonds that are significantly shorter (>0.3 Å) in the S536R structure compared to WT are shown as orange and red bars. The majority of these are in domains 1 (green) and 3 (magenta). **B**. Principal component analysis showing PC1 histogram for the ES complex (blue) and E-TS complex (red). **C**. Two-dimensional plot of PC1 versus PC2 for the ES complex (blue) and the E-TS complex (red). Note the similarity between the two areas when compared to Figure 4D giving a calculated value for 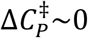.

To check whether these shortened H-bonds are simply a function of the higher resolution of the S566R structure, we refined the S536R structure after truncating the diffraction data to a resolution equivalent to the WT structure (2.30 Å). We repeated the comparison and this also showed a significant shortening of 26 H-bonds at this resolution, confirming that the analysis was not biased based on the very high resolution of the S536R structure.

We carried out extensive MD simulations (20 trajectories of 500 ns each) for the S536R mutant using this high-resolution structure with substrate (ES) or transition state analogue (E-TS) bound to calculate 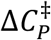 for this mutant. The calculated values for 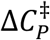 for this mutant are close to zero for all window sizes. This implies that S536R MalL does not fully escape the energetic basin of the transition-state like conformation, either with the substrate or the transition state analogue. PC analysis indicates that conformational sampling of the substrate-complex is indeed more restricted than for WT MalL, with the main conformation being similar to that found with the transition state analogue (Figure 5B&C).

The very high resolution of the crystal structure combined with the reduced conformational dynamics over the time course of the simulations suggests that this conformation is similar to that of the TLC for MalL.

## Discussion

### Interpretation of kinetics data and analysis

There are several possible interpretations to account for the two state model that we present here (MMRT-2S). There is clearly a transition between low temperature behaviour and high temperature behaviour (independent of denaturation). We propose that this transition is cooperative and two-state. The simplest explanation for the low temperature behaviour is that this constitutes the chemical step from TLC to products and that the TLC is favoured at low temperatures (see Scheme 1), as we would expect for the more ordered state (when TLC is compared to ES). At high temperatures, the activation barrier Δ*G*^‡^ involves a large, negative 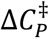 value and steeply temperature-dependent values for Δ*H*^‡^ and Δ*S*^‡^. The fact that Δ*H*^‡^ proceeds from positive to negative values suggests that this step involves a cooperative conformational process consistent with the ES-to-TLC transition. Whether there is a change in rate determining step or whether the conformational transition is combined with the chemical step (c.f. the equilibrium model) is not clear. Nonetheless, the emergence of negative activation enthalpies under either the equilibrium model or a change in rate determining step implies that the conformational transition plays a significant role in the observed kinetics at these intermediate and high temperatures.

### Relationship to other models

The equilibrium model postulates two conformations, one of which is inactive (*E*_*act*_ and *E*_*inact*_).^32^ This is consistent with the ES and TLC conformations postulated here. If we assume that the ES state is saturated, and that *k*_*chem*_ is rate limiting at all temperatures, then the rate equation simplifies to:

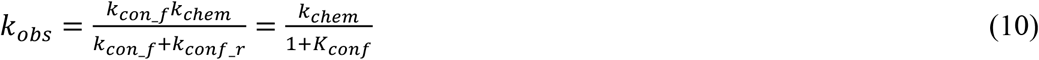

Under conditions where the conformational equilibrium, *K*_*conf*_, is very small (in the case of MalL, at low temperatures) the rate is primarily a function of the chemical step. When the temperature increases, the rate is modulated by the equilibrium between the ES (*E*_*inact*_) and TLC (*E*_*act*_) states. Thus, at high temperatures the activation barrier is a combination of conformational changes and the chemical step. As we have determined that there is a significant 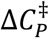 for the equilibrium between ES and TLC, then the expression for Δ*G*^‡^ for the equilibrium model becomes:

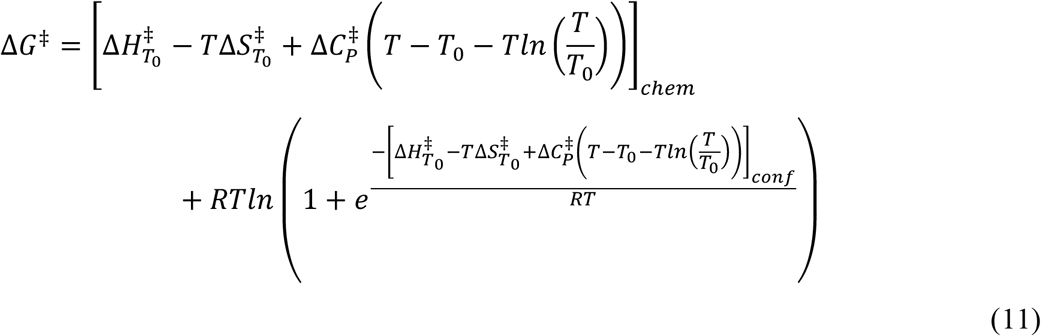

…where the first term in square brackets refers to the chemical step (with 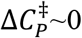) and the exponential term refers to the conformational step (with 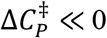). This model fits the data very well (final line in Table 1, Figure 6) although the fitting errors are large as we would expect from fitting 6 parameters. Further problems with fitting also arise from the exponential term making this model very difficult to unambiguously determine. Nonetheless, there are significant differences in the predicted temperature-dependence of the activation heat capacity when comparing our two-state model (MMRT-2S) and the equilibrium model. It is not possible to discriminate between the equilibrium model and the two-state model given the data (Figure 6A&B, the sum of squares for the residuals are similar for the two fits). If the observed changes were genuinely due to a change in the rate determining step from the chemical step to the conformational step then a suitable test would be to find mutants that showed an increase in rate at high temperatures where *k*_*conf_r*_ becomes significant and thus, show that the conformational step is rate limiting at high temperatures.

**Figure 6.**
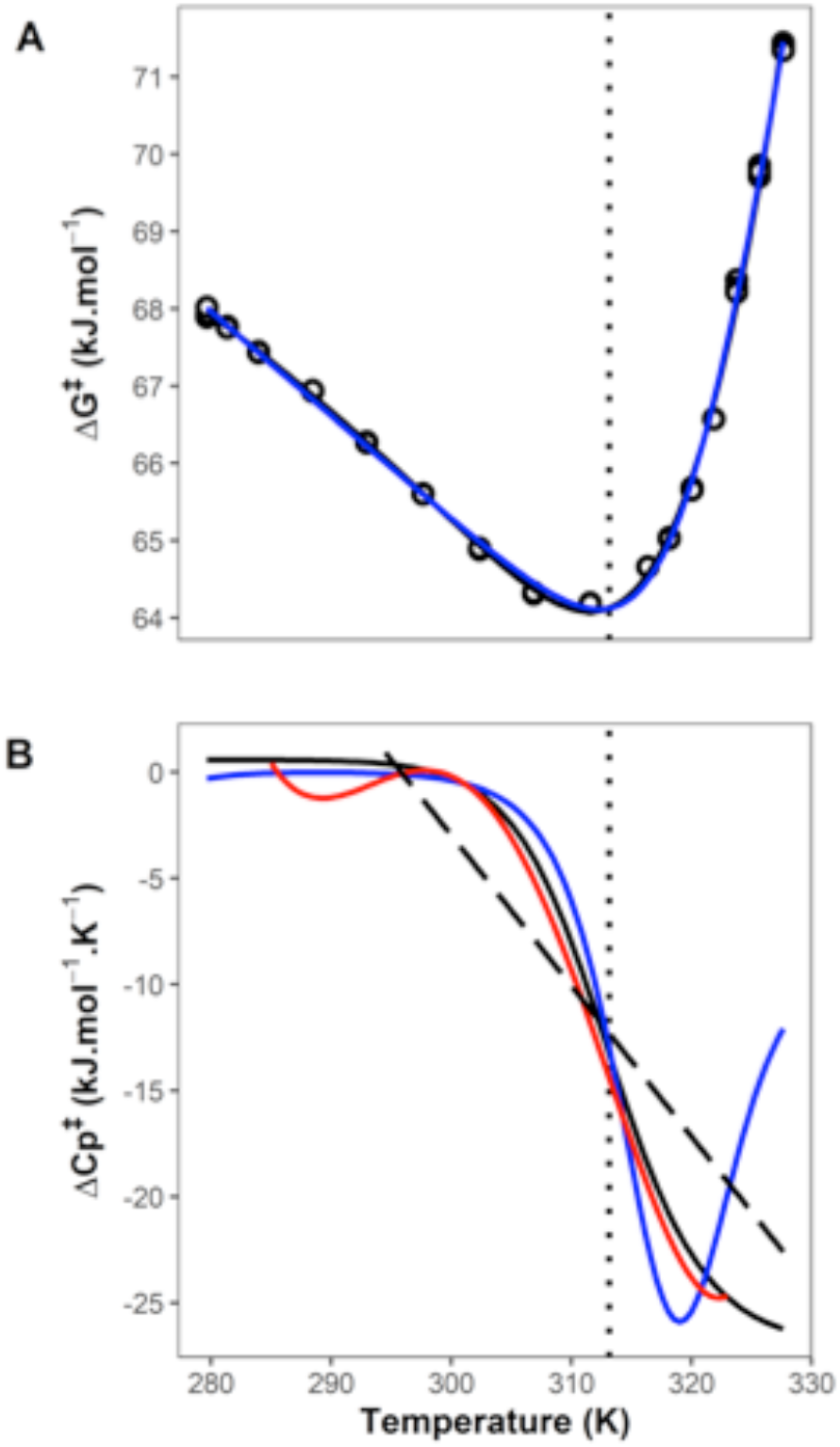
Comparison between two-state (MMRT-2S) and equilibrium models. **A**. Δ*G*^‡^ versus T fitted using MMRT-2S (black) and equilibrium model (blue). **B**. 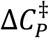 versus T derived from the fits in A. Black is the MMRT-2S, dashed black line is MMRT-1L (i.e. linear 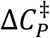, equation 6), blue is the equilibrium model (equation 11) and red is derived from a polynomial fit of order 6 to Δ*G*^‡^ and calculating 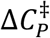 from the second derivative.

In their analyses, Åqvist and colleagues have suggested that the “correct” model in the case of α-amylase AHA and MalL is an equilibrium model with an off-pathway dead-end conformation and that the chemical step is rate limiting at all temperatures.^5^ However, they present Δ*H*^‡^ as being large and positive at low temperatures and then large and negative at high temperatures. It is difficult to imagine a chemical step where this is the case without contributions to the activation barrier coming from a conformational change. It is also hard to rationalise the presence of an off pathway conformation from an evolutionary perspective. These issues are resolved if, instead, the equilibrium is on-pathway between ES and TLC conformations and involves a cooperative conformational transition contributing to the activation barrier.

The combination of high-resolution temperature-dependent enzyme kinetics, molecular dynamics simulations and X-ray crystal structures provide evidence for an on-pathway equilibrium between ES and TLC conformations for MalL. This makes intuitive sense insofar as the enzyme-substrate complex must visit a conformation that favours the chemical transition state species in order to facilitate catalysis. If the on-pathway equilibrium involves a cooperative transition, then we would expect a change in heat capacity for this equilibrium. A proxy for this ES-TLC equilibrium can be found in the binding of a transition state analogue to the enzyme MTAP and this binding is accompanied by a large negative value of Δ*C*_*P*_ as measured by isothermal titration calorimetry (^−^2.4 kJ.mol^-1^.K^-1^).^13^ In this case, the enthalpy of binding is large and positive at low temperatures (∼20 kJ.mol^-1^ at 293 K) and large and negative at high temperatures (∼^−^40 kJ.mol^-1^ at 319 K). Depending on the position of the transition state for the equilibrium, we would also expect a change in activation heat capacity for the kinetics of this transition. In the case of MTAP, curvature in the observed temperature-dependence of the kinetics for this enzyme give very similar values for the activation heat capacity, 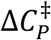 (^-^2.3 kJ.mol^-1^.K^-1^). The equilibrium constant for the interaction between MTAP and various transition state analogues is in the range ∼10^−9^ – 10^−10^ M. By comparison, in the case of glycosidases, we are expecting an equilibrium constant for transition state binding of ∼10^−19^ M based on the rate enhancement over that in water at pH 7.0.^33,34^ Thus, we might expect larger absolute values of Δ*C*_*P*_ for the ES-TLC equilibrium and by implication, the *kinetics* of the ES-TLC transition will also occur with 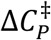 depending on the position of the transition state with respect to the ES and TLC conformations. This will entail a curved temperature-dependence of Δ*G*^*‡*^. Notably, the kinetics of cooperative processes are often slow (akin to folding kinetics) and thus, may be on a similar timescale to the chemical step for catalysis.

Conceptually, the ES-to-TLC transition can be considered under three potential regimes. The first is if the barrier between ES and TLC is very low at all temperatures (compared to the chemical step). In this case the ES/TLC can be considered as one species that is simply conformationally dynamic. Here, the equilibrium model is valid (equation 10), the TLC is always accessible and the rate is simply the chemical step with expected Arrhenius-like behaviour. The second scenario is that there is a moderate barrier between ES and TLC and that this barrier is temperature-dependent. In this case the observed rate will be a combination of the chemical step and the conformational step and that this will be more marked at low or high temperatures. This is the case in our analysis of MalL, and the analysis of Warshel for ADH, giving significant 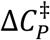 values at high and low temperatures respectively. When the ES state is favoured, the free energy landscape is broad and must proceed through a TLC bottleneck to reach the chemical step giving rise to large negative values for 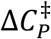 and this could occur at low temperatures (e.g. ADH) or high temperatures (e.g. MalL) (see Figure 7). The third scenario is that the ES-TLC barrier is extremely temperature-dependent and thus is negligible at one temperature and dominates at another. This constitutes a genuine change in the rate determining step. It is difficult to distinguish between the latter two scenarios and this requires further investigation. An obvious route here would be to find mutations that change the conformational barrier which would alter the temperature-dependence of the rate in the case where there has been change in rate determining step.

**Figure 7.**
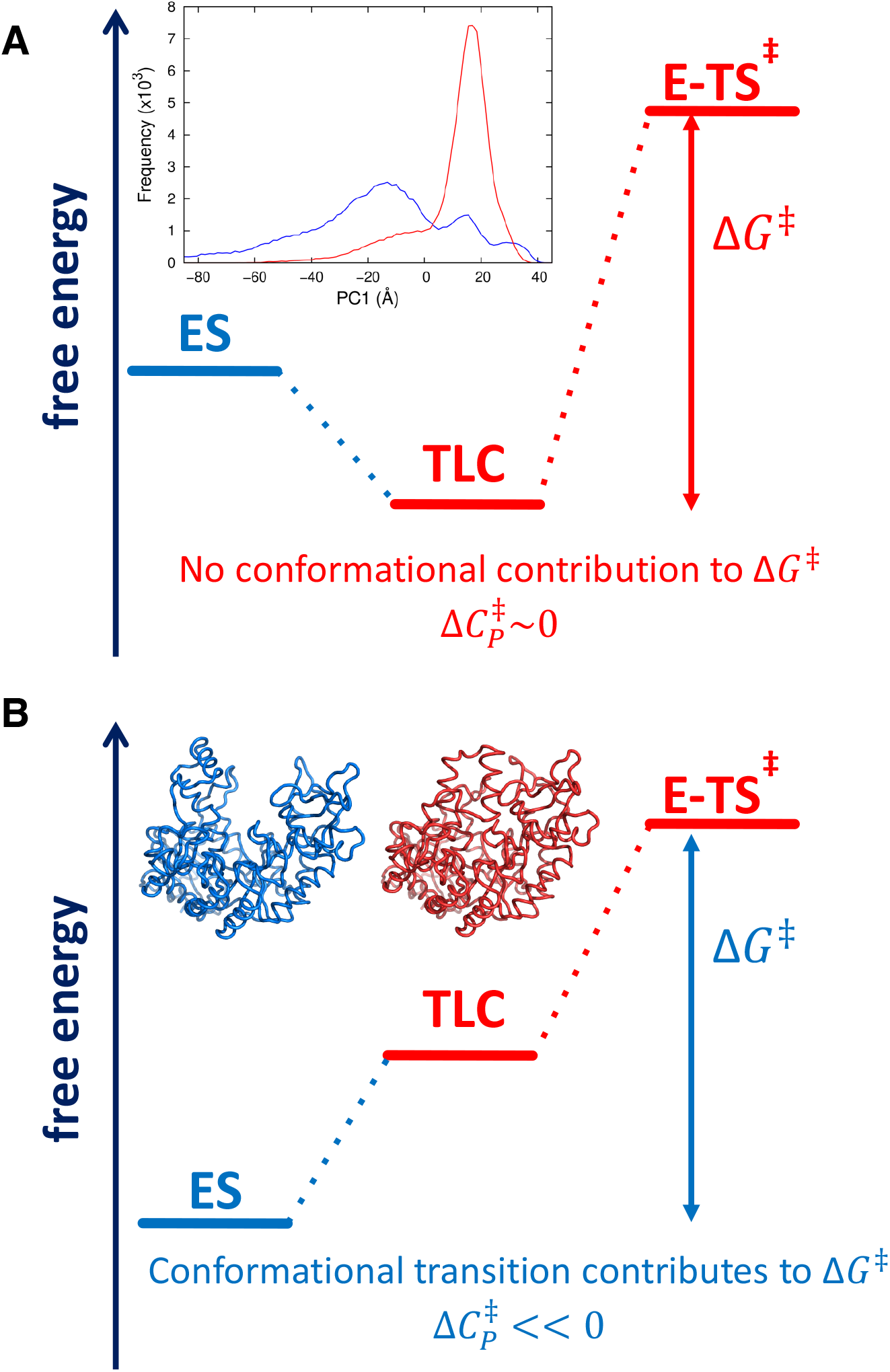
Free energy schematic at two different temperatures. **A**. If the temperature favours the TLC, then there is no contribution from conformational changes to Δ*G*^‡^ and 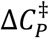 will be near 0 (i.e. the heat capacity for E-TS^‡^ is similar to that for TLC). This is the case for MaL at low temperatures and ADH at high temperatures. **B**. If the temperature favours the ES state, then conformational fluctuations will be a component of Δ*G*^‡^ and 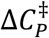 will be significantly less than 0. This is the case for MaL at high temperatures and ADH at low temperatures.

Our two-state model (MMRT-2S) describes this process well. Its interpretation is consistent with the conformational change as the main contributor to 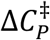 irrespective of the microscopic details of the mechanism. We present scheme 1 as a general scheme for enzymes: the role of an enzyme is to bind the substrate relatively weakly and the transition state much more strongly.^35,36^ The scheme is a minimal model. The reality of enzyme conformational behaviour is of course probably much more complicated, with many possible conformations distinguishable in various ways. Nonetheless, the ES conformation must visit a conformation that facilitates catalysis (i.e. the TLC) and this is discussed further below. This argument is consistent with previous arguments that highlight the role of a specific conformation of the enzyme which lowers the chemical reaction barrier.^37,38^ Our argument is that forming this conformation involves a cooperative transition to instigate the very large apparent *K*_*M*_ values for the transition state (i.e. precise preorganisation). This can be seen in the extensive molecular dynamics simulations of the ES complex and the E-TS complex, for which principal component analysis shows that the ES complex visits the TLC and the TLC is much more constrained in comparison to ES (Figure 4).

MMRT-2S is rather complex to fit without a large number of data points. Much of the temperature-rate data in the literature contain relatively few points and for such typical cases, we recommend using a linear 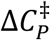 model as a convenient approximation (MMRT-1L). This linear model clearly reveals details of the ES-TLC equilibrium and temperatures at which the conformational component contributes significantly to the activation free energy. In the case of ADH, this occurs at low temperatures (with a positive slope for 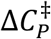, Figure 3) and in the case of MalL this occurs at higher temperatures (with a negative slope for 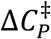, Figure 2). Indeed, this approach may be used to glean evidence for the nature of the TLC conformation as the steeper the slope for 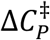, the greater the cooperativity of the transition.

Our two-state model (MMRT-2S) is consistent with the numerous historical arguments that postulate two states for enzymes with and without allosteric regulation. Here, we extend these arguments to identify and characterise the two states, ES and TLC. Our arguments are consistent with those of Åqvist and colleagues (i.e. the equilibrium model), except that the equilibrium is on path for ES and TLC. In the case of the directed evolution of the activity of a designer Kemp eliminase enzyme, which gives rise to curved Arrhenius plots, we argue that this is most obviously the result of the optimisation of the TLC to improve preorganisation and catalysis with increases in ES-TLC cooperativity and attendant correlated motions at the TLC.^30,39^ The increases in cooperativity to reach the TLC also rationalise remote mutations that improve catalysis as seen in many directed evolution studies.^40,41^

### Evidence for TLCs in other enzymes

Molecular simulations show that specific reactive conformations of enzyme-substrate complexes are involved in many enzymes, and identify structural features of these complexes, providing details of TLCs. One example is fatty acid amide hydrolase, in which hydrolysis of oleamide occurs by a distinct, high energy conformation: the barrier to reaction is significantly lower in this conformation, and so reaction will proceed via this TLC, even though it is much less populated than other conformations of the ES complex of FAAH at 300K.^42,43^ In lactate dehydrogenase, while reaction in the direction of lactate formation can proceed with the active site loop open or closed, oxidation of lactate to pyruvate requires loop closure.^44^ In thymidylate synthase, different experimentally observed conformations of the enzyme show different reaction barriers, associated with different product stabilization.^45^ From simulations of HIV-1 protease, Ribeiro *et al*. noted that the reaction will be dominated “by a very few transient enzyme conformations that provide very low barriers”.^46^

QM/MM and MD simulations of ketosteroid isomerase (KSI) show that changes in active site structure cause changes in solvation of the catalytic base (Asp38): these changes significantly lower the barrier to reaction; the reaction proceeds via the TLC in which the base is less solvated.^38^ In triosephosphate isomerase (TIM), a crucial active site loop adopts various different conformations in the ES complex, but a low barrier to reaction is only found when the loop is fully closed: in the TLC, the catalytic base (Glu165) is desolvated and so is more basic.^37^ Reactive conformations (TLCs), involving distinct conformations of the substrate and active site, and desolvation of the catalytic base, are also important in the antibiotic breakdown activity of β-lactamases.^47,48^ Mhashal *et al*. showed that reaction in glycerol-3-phosphate dehydrogenase proceeds via a conformation in which the active site is also less solvated.^49^ Changes in basicity associated with changes in solvation, and active site loop behaviour, have also been found to be important in evolution and activity of dihydrofolate reductase^50^, and in differences in reactivity between thermophilic and mesophilic DHFR, which are also modulated by dynamical/entropic changes caused by dimerization of the thermophilic enzyme.^51^ Changes in solvation (e.g. desolvation of catalytic carboxylate groups) are likely to be a common feature of TLCs. In the TLC, specific changes in solvation increase reactivity of important groups, achieving desolvation and ground state destabilization within the overall context of a polar active site.

It is important to note that a TLC is a distinct conformation of the enzyme-substrate complex, involving structural changes throughout the protein, and is not limited to trivial substrate conformational changes or simple loop opening and closing motions. The conformation of the protein as a whole is different. The dynamics of different parts of the protein change in different ways in the TLC: some regions becoming less ordered, and others becoming more flexible. This is shown by MD simulations of KSI: in this homodimeric enzyme, the dynamics of the monomer in which reaction is not occurring are strongly affected by changes in the reactive monomer.^14^ As noted above, the dynamics of the small domain of MalL, far from the active site, are different in the TLC. The relative stabilities of TLCs may be modulated by evolution, allosteric ligands and solvent effects as well as by temperature^52^

## Conclusions

Warshel and colleagues have suggested that the abrupt changes seen for the temperature-dependence of Δ*H*^‡^and Δ*S*^‡^ for ADH are indicative of a phase transition.^8^ This description is useful because the thermodynamic tools used to describe phase transitions are very well studied.^53^ Indeed, the two-state model that we present here is the analogue of a second order, finite-size phase transition.^54^ In this case, the kinetics of the system are fundamentally altered from when the temperature favours the TLC state (e.g. MalL at low *T*) to when ES state is favoured (e.g. MalL at high *T*). The kinetics of phase transitions have also been very well studied and are generally described in terms of a sphere of nucleation which must reach a critical size in order to affect the phase transition. Similar hypotheses have been put forward for protein folding (e.g. the nucleation-condensation model^55^) and may be a feature of the ES- to-TLC transition for enzyme-catalysed reactions.

Here, high resolution temperature-rate data provide sufficient detail to characterise a two-state cooperative conformational transition prior to the chemical step for enzyme catalysis (Figure 7). From high resolution structures and molecular dynamics simulations, we have characterised these two states (ES and TLC). At temperatures that favour the TLC (Figure 7A), enzyme kinetics are a function of the chemical step (e.g. low temperatures for MalL and high temperatures for ADH). In contrast, at temperatures that favour the ES (Figure 7B), the enzyme kinetics are a combination of both a conformational step and a chemical step leading to significant negative values of the activation heat capacity 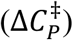 at these temperatures (high temperatures for MalL and low temperatures for ADH). Finally, a model that uses a linear change in 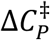 (MMRT-1S) as an approximation can discriminate between MalL-like behaviour and ADH-like behaviour and is sufficiently simple to use with most published data. This approach should find wide application in analysing and understanding the temperature dependence of enzyme-catalysed reactions.

## Methods

### Protein Expression

*Bacillus subtilis* MalL, and single amino acid variants, were expressed with N-terminal hexa-histidine tags in *E. coli* BL21 DE3 cells. Luria Broth cultures in exponential phase were induced at 18 °C with 0.75 mM Isopropyl β-D-1-thiogalactopyranoside and grown overnight.

### Protein Purification

Protein purification was carried out in two steps via immobilised metal affinity chromatography and size exclusion chromatography (IMAC and SEC respectively) at pH 7.0. Initially, cell pellets were lysed via sonication on ice. A 25 mM to 0.5 M imidazole gradient over 50 mL was used to elute MalL during IMAC. SEC was carried out in 20 mM HEPES buffer. Enzymes were dialysed into 40 mM NaPO_4_ buffer with 150 mM NaCl.

### X-ray Structure Determination

Crystallisation of MalL S536R was performed using hanging-drop vapour diffusion at 18 °C. Crystals were obtained in 0.1 M Tris pH 8.0, 0.2 M ammonium acetate and 18% w/v PEG 10,000. Data collection was performed on flash-cooled crystals on the MX2 beamline at the Australian synchrotron. A solution of 0.1M Tris pH 8.0, 0.2 M ammonium acetate and 17% w/v PEG 10,000 with 20% v/v glycerol was used as cryoprotectant. Data was indexed, integrated and scaled in XDS^56^ and further scaled and merged in Aimless.^57^ The structure was solved by molecular replacement in Molrep^58^ with wildtype MalL (PDB code: 4M56) as the search model. This was followed by iterative cycles of manual building in COOT,^59^ structure correction using PDB-REDO,^60^ and further refinement using Phenix.Refine^61^ and Refmac5.^62^

### MalL Temperature Assay

The KinetAsyst™ Stopped-Flow System (TgK Scientific, UK) with a connected circulating water bath for temperature control was used to characterise the temperature profiles of MalL via cleavage of saturating concentrations of *p*-nitrophenyl-α-D-glucopyranoside at 405 nm. Reactions were completed in triplicate with five 0.2 s dummy shots in between. Each reaction was carried out for 45 s. Temperature values reported are those from the thermostat control monitoring the reaction chamber. Enzyme stability over the experimental time-period (∼5 hours) was confirmed by a mid-range temperature assay at the end of the experimental time-period.

### Rate Calculation

Linear regression of (at most) the first 10 s of the reaction was carried out using Kinetic Studio (TgK Scientific, UK). Catalytic rates (*k*_*cat*_; s^-1^) were determined using an extinction coefficient (L mol^-1^ cm^-1^) of 7413. Rate data were converted to change in Gibbs free energy (*ΔG*^‡^) for model fitting using the Eyring equation with transmission coefficient set to 1.

### Model Fitting

Levenberg-Marquardt non-linear regression was carried out in RStudio.

### Molecular Dynamics Simulations and Principal Component Analysis

We previously performed 10, 500 ns long molecular dynamics (MD) simulations of WT-MalL with both isomaltose bound (reactant state, RS) and with a transition state analogue bound (TSA).^14^ Here, we performed an additional 10 MD simulations of WT-MalL alongside 20 new MD simulations of the point variant S536R (with the starting coordinates based on the new X-ray structure), in both their ES and E-TS states, meaning each state was sampled with twenty 500 ns replicas each. The same protocols, force-field parameters (ff99SB-ILDN for protein, TIP4P-Ew for water, GLYCAM 06j-1 for isomaltose and combination of GLYCAM 06j-1 and GAFF for the TSA) and protonation states were used as in our previous work.^14^

Trajectory analysis was performed using CPPTRAJ (part of the AmberTools suite of programmes https://ambermd.org/AmberTools.php) using snapshots taken every 10 ps from all trajectories unless otherwise stated. The Cα RMSFs were determined by RMSD fitting (to the Cα of residues 7-561) to a running average coordinates using a time window of 10 ns. A hydrogen bond (HB) was defined to exist using typical criteria if the donor-acceptor distance was within 3.5 Å, and if the donor-hydrogen-acceptor angle was within 180±45°,. Hydrogen bonds between all residues were separated into main chain (MC) and side chain (SC) contributions from each residue, giving rise to either MC-MC, MC-SC and/or SC-SC HBs between residues. The average difference (from the 20 replicas) between the RS and TSA was determined from the 20 runs, and the significance of the differences was evaluated using a *t*-test. Principal component analysis (PCA) was performed on the Cα of every residue for all states simulated (WT, and S536R in both RS and TSA forms) combined. RMS fitting was first performed to a crystal structure of WT-MalL (PDB: 5WCZ) using the Cα of residues 7-561, to create an average structure. Following this, all snapshots were then re-fitted to this average structure for the subsequent calculation (again to the Cα of residues 7-561).

### Hydrogen bond analysis

Hydrogen bonds were analysed for WT MalL and S536R structures. All hydrogen bonds were found using FindHBond in Chimera 1.15.^63^ Hydrogen bond criteria are described in Mills & Dean (1996)^64^ with criteria relaxed by 0.4 Å and 20°. Existing explicit hydrogens were removed from the structure prior to analysis. Unique H-bonds and those that were at least 0.3 Å shorter than their equivalent were extracted. To discount the effects of the resolution the structures were solved at (WT: 2.3 Å, S536R: 1.1 Å) the process was repeated with the S536R structure where the data were truncated to 2.3 Å resolution and the structure refined. In addition, bonds involving multiple rotamers, or in regions not modelled in the other structure, were discounted.

## Acknowledgements

This research was undertaken in part using the MX2 beamline at the Australian Synchrotron, part of ANSTO, and made use of the Australian Cancer Research Foundation (ACRF) detector. VLA, LS & AJM are investigators funded by a Marsden Fund Council grant from the Marsden Fund of New Zealand. EJW, CJH and AW acknowledge doctoral funding from the University of Waikato. M.C. and A.J.M. thank the EPSRC Centre for Doctoral Training in Theory and Modelling in Chemical Sciences (EP/L015722/1). We further acknowledge EPSRC funding for CCP-BioSim (EP/M022609/1). This work is part of a project that has received funding from the European Research Council under the European Horizon 2020 research and innovation programme (PREDACTED Advanced Grant Agreement no. 101021207) to A.J.M. The simulation work was conducted using the computational facilities of the Advanced Computing Research Centre, University of Bristol. MWvdK thanks BBSRC for funding (BB/M026280/1).

## Supplementary Material

**Figure S1.**
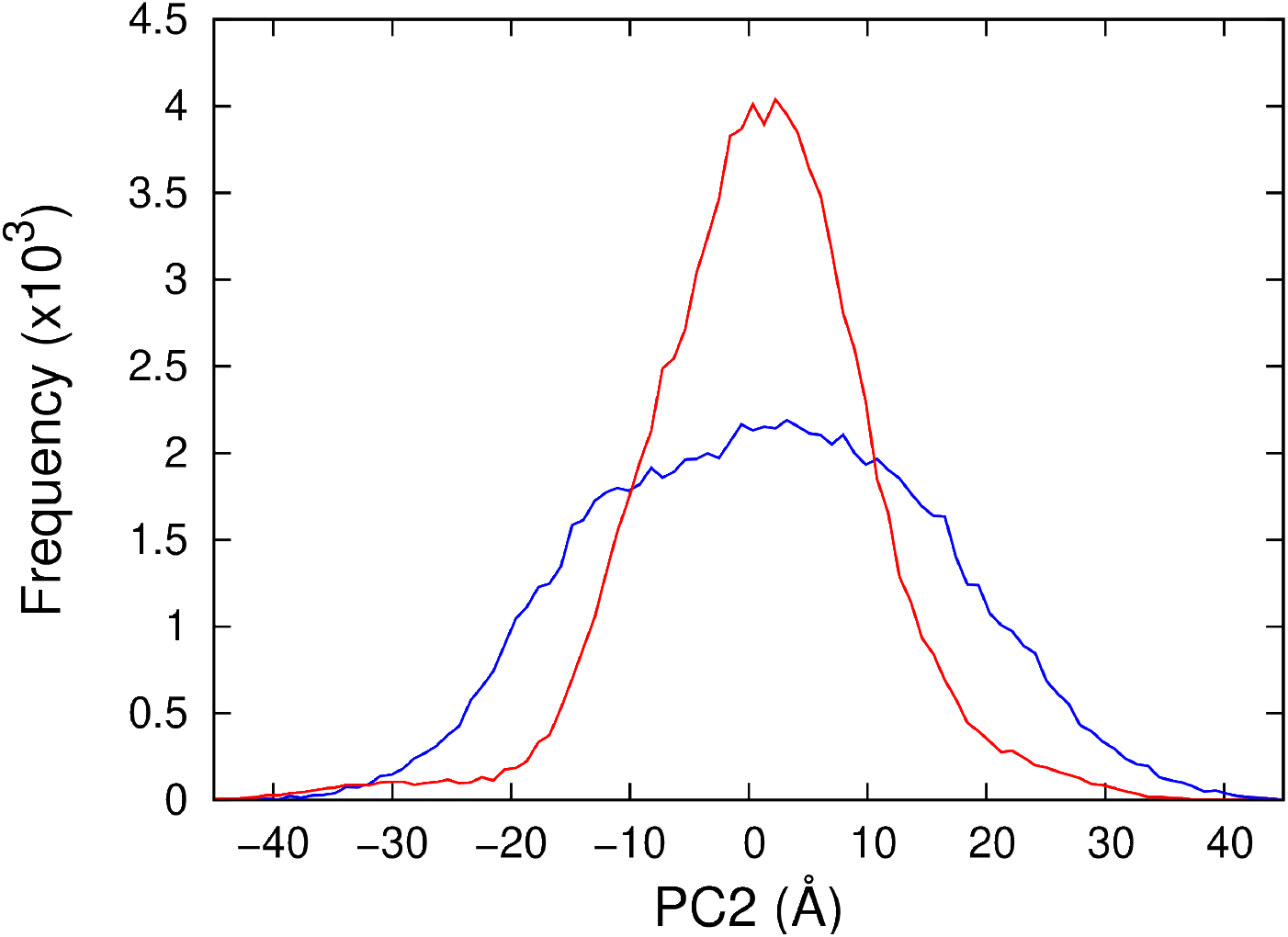
Principal component analysis for WT MalL in the ES and E-TS states showing a projection of the second principal component, PC2.

**Figure S2.**
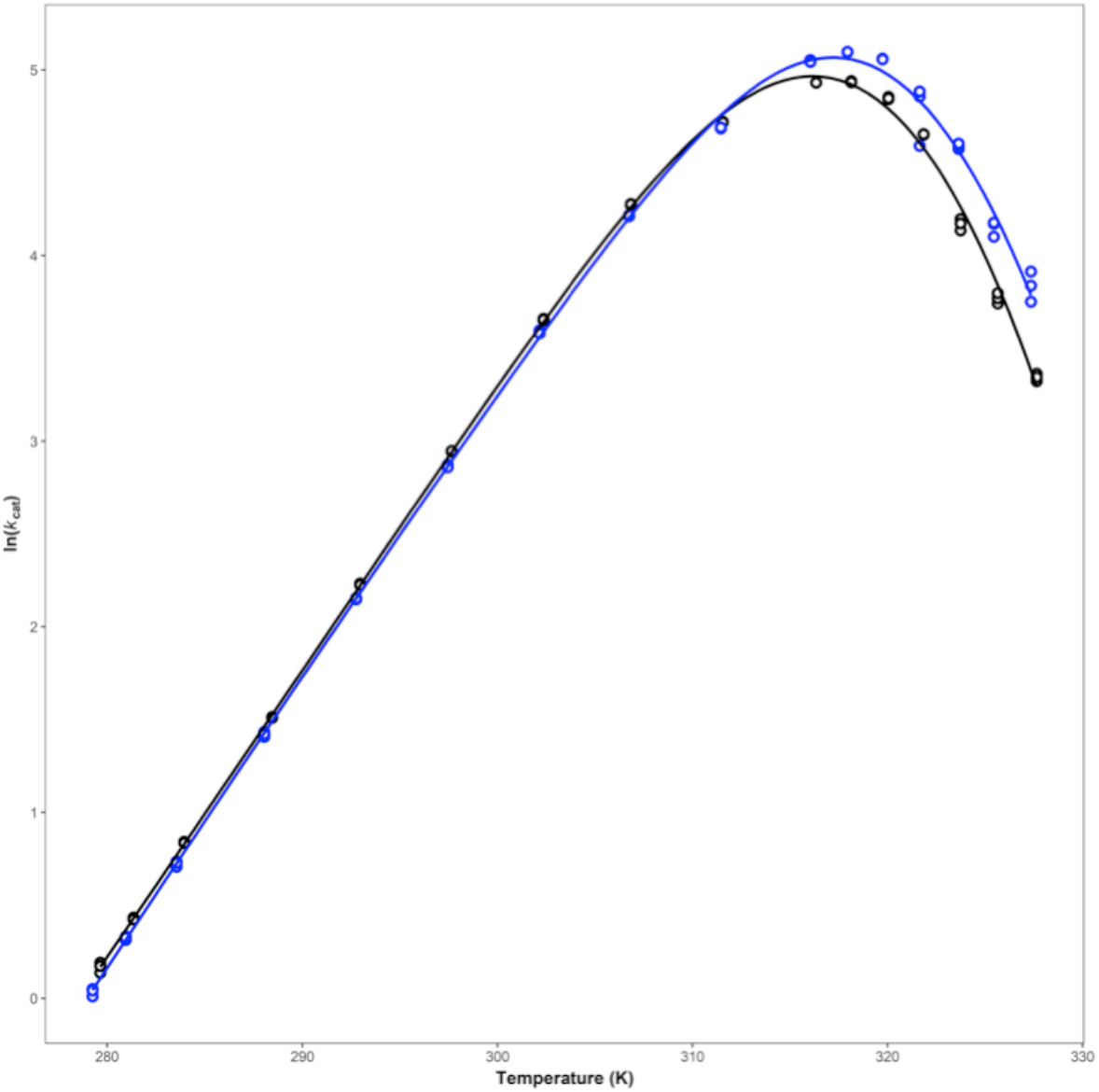
Ln(rate) versus temperature for WT MalL (black) and S536R (blue).

**Table S1.** Data collection and refinement statistics for MalL S536R.

| Statistic | Mall S536R |
| --- | --- |
| Wavelength (Å) | 0.953735 |
| Space group | P 1 2<sub>1</sub> 1 |
| Unit cell lengths (Å) | a = 48.75 b = 101.00 c = 61.75 |
| Unit cell angles (°) | $\alpha$ = 90.00 $\beta$ = 113.06 $\gamma$ = 90.00 |
| Resolution (Å) | 1.10 - 44.85 (1.10 - 1.12) |
| R<sub>merge</sub> | 0.107 (0.548) |
| Completeness (%) | 94.2 (87.1) |
| Redundancy | 10.9 (6.9) |
| No. of observations | 2270037 (65265) |
| No. of unique reflections | 208774 (9509) |
| Mean I/σI | 12.7 (2.9) |
| R factor | 0.126 |
| R<sub>free</sub> | 0.145 |
| Protein atoms | 9633 |
| Solvent atoms | 785 |
| Average temperature factor (Å<sup>2</sup>) | 16.13 |
| RMSD bond lengths (°) | 0.01 |
| RMSD bond angles (Å) | 1.182 |

